# Higher gene expression variability in the more aggressive subtype of chronic lymphocytic leukemia

**DOI:** 10.1101/005637

**Authors:** Simone Ecker, Vera Pancaldi, Daniel Rico, Alfonso Valencia

## Abstract

**Background:** Chronic Lymphocytic Leukemia (CLL) presents two subtypes which have drastically different clinical outcomes. So far, these two subtypes are not associated to clear differences in gene expression profiles. Interestingly, recent results have highlighted important roles for heterogeneity, both at the genetic and at the epigenetic level in CLL progression.

**Results:** We propose to use gene expression variability across patients to investigate differences between the two CLL subtypes. We find that the most aggressive type of this disease shows higher variability of gene expression across patients and we elaborate on this observation to produce a method that classifies patients into clinical subtypes. Finally, we find that, overall, genes that show higher variability in the aggressive subtype are related to cell cycle, development and inter-cellular communication, probably related to faster progression of this disease subtype.

**Conclusions:** There are strong relations between disease subtype and gene expression variability linking significantly increased expression variability to phenotypes such as aggressiveness and resistance to therapy in CLL.

## Background

One of the outstanding challenges in biology is elucidating the relationship between genome, epigenome and phenotype. Notwithstanding the considerable progress that has been made in terms of mapping the epigenetic state of cells along with their transcriptome, it has often been hard to see the interdependencies between the two and their joint contribution to cellular behavior. We are just starting to unravel the different genetic and non-genetic factors that are responsible for the incredible variability of phenotypes that can be observed in a population of cells.

Biological noise is emerging as an important factor influencing the phenotypic variability in cell populations. The first experiments measuring fluorescence of reporters in single bacteria [1] highlighted the presence of various sources of ‘noise’ that would contribute to the variability observed. Intrinsic noise, which is inherently caused by stochasticity in the biochemical phenomena that lead to gene transcription and affects each gene independently, and extrinsic noise, which causes fluctuations in the value of expression correlated amongst multiple genes [1]. In fact, biological phenomena are governed by randomness just like other physical systems on the small scale. For example, the production of mRNA happens in bursts whose regulation in size and frequency can control not only the average amount of RNA produced, but also the fluctuations in this value [2].

Recently, single-cell methods in yeast and mammalian systems have studied noise and cell-to-cell variability, which is now recognized to be at the basis of many interesting biological processes, for example *p53* oscillations [3] and *NF-κB* pulses of localization in the nucleus [4, 5]. The gene expression variability at the single cell level is probably having an effect on the variability across different organisms in a population. Indeed, a strong correspondence between expression variability due to stochastic processes in single cells from the same population and variability of gene expression in a population measured across different conditions is commonly observed. Multiple experimental investigations of this relationship have led to accept that common mechanisms are probably responsible for the two different types of variability of gene expression, connecting variability in a population to variability across a time courses [6, 7]. The conclusion from these studies is that variability across conditions in a time course, between different individuals that have slightly different genetic backgrounds, and variability in single cells of the same isogenic population are strongly related. This allows us to measure variability of one type and use it as an estimate of the other types of variability.

It is therefore fair to ask what regulates the weaker or stronger propensity of a gene to be regulated, both in terms of plasticity in different conditions and in terms of stochastic noise. It is widely recognized that specific promoter structures (TATA boxes) are found mainly in genes with functions related to the response to external stimuli, which are also genes that usually have widely fluctuating single-cell levels within populations [8–10]. The characteristics and dynamics of regulation are very likely related to the chromatin structure in the region of the promoter of the gene and, more specifically, to the nucleosome distribution [11].

This biological observation is reminiscent of a widely accepted concept in physics which goes under the name of ‘fluctuation dissipation theorem’ and states that quantities that are observed to stochastically fluctuate on a large scale are also likely to have large responses to a stimulus, whereas quantities that have limited stochastic fluctuations will have smaller responses to the same size stimulus [12]. The analogy with gene expression would suggest that when a gene needs to undergo large changes in its levels in response to signaling, for example, it will be easier to achieve that level if the gene already displays large stochastic fluctuations in the absence of the stimulus.

It is well known that tumors show increased heterogeneity compared to normal tissue [13–15]. The presence of heterogeneity in tumors is furthermore known to affect aggressiveness and resistance to therapy [16, 17], but is traditionally investigated in solid malignancies, which can present a very diverse population of clones. However, even hematologic diseases, which are thought to arise from clonal populations, can display a degree of genetic and non-genetic heterogeneity [18].

In this work we will focus on gene expression variability between individuals. Variability of gene expression has been suggested to be an important parameter to be measured alongside the average levels of gene expression [19, 20]. We focus on two datasets of chronic lymphocytic leukemia (CLL) – a B-cell neoplasm – in which gene expression was measured for large cohorts of patients in two independent datasets [21, 22] and for which clinical data was also available. Two major subtypes in CLL are defined by the mutational status of the immunoglobulin heavy chain variable region (IgVH).

CLL patients showing fewer mutations in this region, defined as U-CLL (‘unmutated’ CLL), have a worse prognosis compared to M-CLL (‘mutated’ CLL) patients, who show a larger number of mutations in the IgVH gene region [23]. The study of the International Cancer Genome Consortium (ICGC) [22] showed that, although there are significant differences in the methylome of M-CLL and U-CLL, a strong correspondence of DNA methylation with gene expression levels was not found. A subsequent study of the ICGC [24], which extensively characterized the transcriptome of CLL using RNA sequencing, revealed two new subtypes of the disease, which are completely independent of the well characterized clinical subgroups based on the mutational status of the IgVH. This further demonstrated that the two important clinical subtypes M-CLL and U-CLL do not seem to be directly reflected by gene expression levels.

Interestingly, analyzing the data provided by these two previous studies, we find significant differences in the variability of gene expression between M-CLL and U-CLL. The more aggressive U-CLL subtype exhibits increased expression variability. Even more strikingly, we show that patients can be correctly classified into the two disease subtypes by a machine learning approach solely based on gene expression variability measurements.

In this work we demonstrate that there are strong relations between disease subtype and gene expression variability, suggesting an impact of gene expression variability on tumor adaptability and aggressiveness in CLL.

## Results and discussion

### Inter-patient gene expression variability differs between the two major clinical subtypes of CLL

To quantify the level of variability of tumor samples in the ICGC Chronic Lymphocytic Leukemia (CLL) patient cohort [22, 24] and a second independent CLL dataset used for the validation of the results [21], we study variability in terms of the Coefficient of Variation (CV). The CV is defined as the ratio between the standard deviation of the variable measured across the patients and its mean. As gene expression variability is dependent on the gene expression levels, we analyze the dependence of the CV on the level of expression of the corresponding genes (see Supplementary Figure 1). The relationship between CV and expression level is interesting and non-trivial. The highest levels of expression variability across patients are observed for genes with intermediate levels of expression and not for genes expressed at high or low levels (Supplementary Figure 1). To understand the origin of this behavior it is important to take the intrinsic stochasticity of biological processes into account. The impact of fluctuations is inversely proportional to the number of elements involved in the system. This is a well-established phenomenon observed in physical systems [25] and well characterized in biology [26]. Indeed, there is a component of the CV that is given by the inverse of the mean of expression as a 1/x dependence on expression levels (Supplementary Figure 1). This dependence reflects the fact that introducing one additional element in a small collection (i.e. an extra copy of mRNA of a lowly expressed gene) will have dramatic consequences. In contrast, an extra copy of a transcript that is in high numbers will not produce a substantial change. Stochastic processes of this kind are likely to not be the sole determinants of the CV. The remaining component of the CV is given by the standard deviation of expression, which has a negative quadratic dependence on the mean of expression (Supplementary Figure 1) showing higher values for intermediate expression classes. These observations highlight the importance of taking gene expression levels into account when evaluating gene expression variability in tumoral cells.

Although the CV is the standard measurement of expression variability in the literature, we also employed an alternative measure which has recently been proposed by Alemu et al. [27]. Alemu’s measure of expression variability (EV) tries to account for the above described relationship between mean expression and variability in a distinct way and to provide a measure of variability which is independent of expression mean (Supplementary Figure 2). We observe a high correlation of the CV and EV in both datasets analyzed (Kulis et al. data [22]: Pearson correlation r = 0.74, p-value < 2.2e-16; Fabris et al. data [21]: Pearson correlation r = 0.87, p-value < 2.2e-16; see Supplementary Figure 3) and take both measures of variability into account in all subsequent analyses. In the following we investigate whether gene expression variability differs in clinical subtypes of CLL and whether this difference could therefore be behind the different aggressiveness of M-CLL and U-CLL.

Consistent with our hypothesis, gene expression variability shows a clear difference between the two subtypes (Figure 1A and B) with higher variability associated to U-CLL, the more aggressive disease. On the contrary, the gene expression levels of U-CLL and M-CLL patients showed very little difference (Figure 1C), in agreement with previous reports [28, 29]. These results suggest that expression variability across patients can be an important factor to discriminate the two disease subtypes, for which the general level of expression will not be discriminatory and for which very few differentially expressed genes have been identified [24, 28].

**Figure 1.**
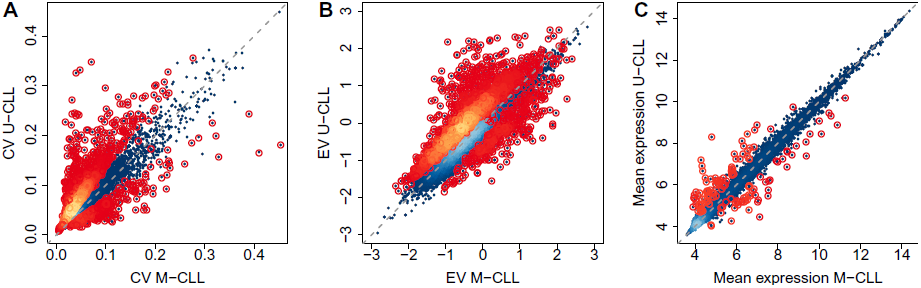
Gene expression variability comparison of M-CLL and U-CLL. Scatterplots comparing M-CLL and U-CLL where each data point represents a single gene. Lighter colors indicate higher densities of data points in the corresponding regions of the plot. Genes with statistically significant p-values at an FDR of 5% are highlighted by circles. The gray dashed line represents the identity line. A) Scatterplot of CV across patients in the two disease subtypes. Genes with statistically significant differential variability according to the F-test (p < 0.05) are highlighted. B) Scatterplot of EV across patients in the two disease subtypes. Genes with statistically significant differential variability according to the F-test (p < 0.05) are highlighted again. C) Scatterplot of mean expression levels across patients in the two disease subtypes. Genes with statistically significant differential expression (|M| >= 1, p < 0.05) are highlighted.

As shown in Figure 1A and B, a substantial number of genes display higher variability across U-CLL patients compared to M-CLL patients. In order to test for statistical significance of these differences, we applied an F-test to compare variances and found 2,025 genes with significantly increased variance in U-CLL whereas only 360 are significantly less variable (FDR = 5%, see Figure 1 and Supplementary Table 1). Repeating this analysis with the dataset of Fabris et al. [21], we confirm the increased variability in U-CLL patients (see Supplementary Figure 4 and Supplementary Table 1) and we see a very strong correlation between the CV of the CLL subtypes in the two patient cohorts (Pearson correlation: M-CLL r = 0.67 and U-CLL r = 0.66, p-values < 2.2e-16, Supplementary Figure 5). Also the differences between CV values for genes in the Fabris et al. [21] and Kulis et al. [22] cohorts are significantly correlated (Pearson r = 0.28, p < 2.2e-16, Supplementary Figure 5) as well as those for the standard deviation (Pearson r = 0.75, p < 2.2e-16, Supplementary Figure 5).). Furthermore, we observe in both datasets a very high correlation of differential variability measured either by CV or EV differences (Kulis et al.: Spearman correlation rho = 0.91; Fabris et al.: Spearman correlation rho = 0.93; all p-values < 2.2e-16, Supplementary Figure 6).

When we take the top 500 genes with increased variability in U-CLL in each dataset (see Additional File X1), we find a significantly higher than expected overlap (69 genes in both lists, Hypergeometric test, p-value < 2.2e-16). Therefore, our results are reproducible in the two datasets, both in terms of i) correlation of the measurements of global expression variability of all genes investigated and in ii) the comparison of ranked lists of the top differentially variable genes. We thus conclude that our findings are unlikely to be caused by batch effects.

We next asked if the differences we observe in expression variability might be explained by differential DNA methylation. For the Kulis et al. dataset DNA methylation data matched to the expression data is available. We therefore compared the methylation profiles of the top 500 differentially variable (DV) genes with increased variability in U-CLL (Additional File X1) but could not observe any strong and clear trend of different methylation levels between the two subgroups investigated (Supplementary Figure 7 and 8).

### Functional analysis of differentially variable genes

We have stated that U-CLL samples exhibit increased expression variability in many genes but we haven’t so far commented on whether specific functional categories are particularly affected by this increased variability of expression. If we assume that most of the differential variability between the two subtypes is due to a biological process, we would expect specific functional classes of genes to be most affected.

Looking at the 500 genes that increase their variability most in the U-CLL patients of the Kulis et al. [22] study (Additional File X1) we observe a very significant enrichment for processes related to the cell cycle, hemopoiesis, multicellular organismal processes, wounding, and an enrichment for development of the immune system and immune system processes (Additional File X2). Performing the same analysis using the Fabris et al. dataset [21] we recapitulate these results to a certain extent, finding significant enrichments in the immune system process, signaling in the immune system and immune response, and – although not reaching statistical significance with an FDR of 5% – in hemopoiesis, development, wounding and cell proliferation (Additional File X2).

To further understand the functional context of these differentially variable genes, we used a B cell specific functional interaction network [30] and extracted a subnetwork of the top differentially variable genes with increased variability in U-CLL in both datasets analyzed (Additional File X1) and their direct neighbors (considering only genes connected with at least two other genes). As a result, we identified a network of 892 genes connected by 3390 edges (see Methods). Figure 2 shows the network with 5 highlighted subnetwork modules we identified (Louvain method [31]). A functional analysis of these network modules shows that every module is highly enriched in biological processes and pathways, further confirming our previous results of biological functions affected by increased expression variability in U-CLL and giving a deeper insight into these processes and pathways as well as the genes involved (Table 1 and Additional File X3).

**Figure 2.**
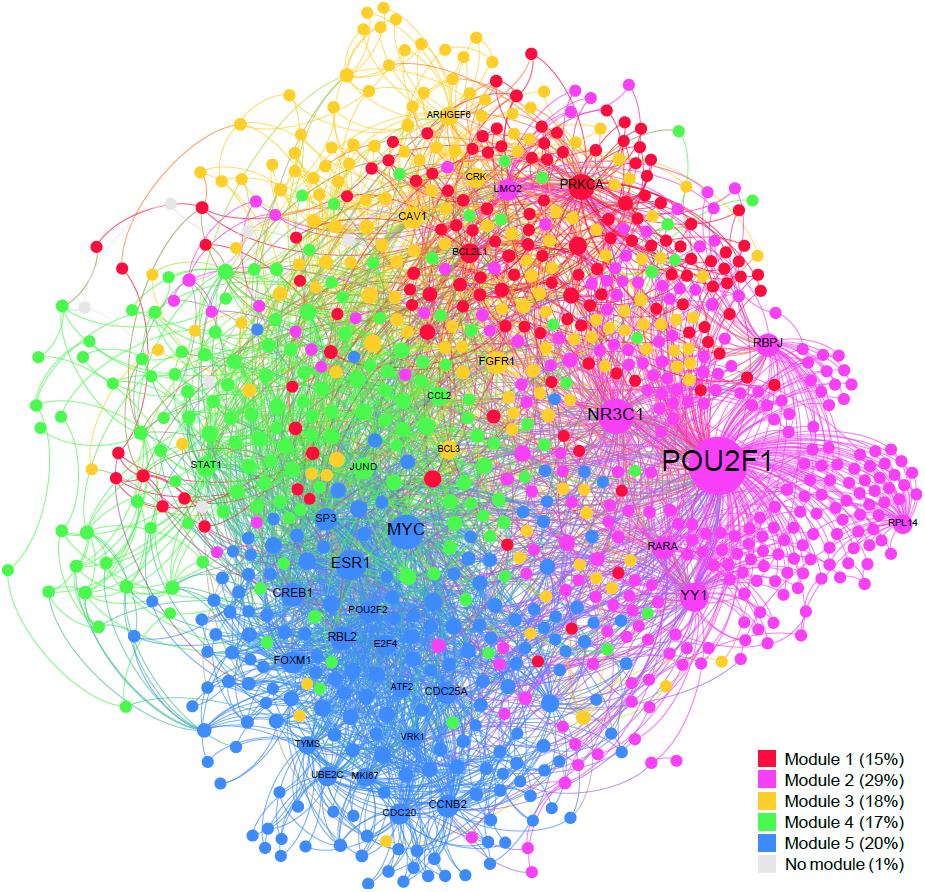
Network representation of genes with increased variability in U-CLL in the context of a B cell specific network [30]. Node sizes are determined by the degrees of the nodes, i.e. big nodes represent highly connected genes. Different modules are highlighted in different colors.

**Table 1.**
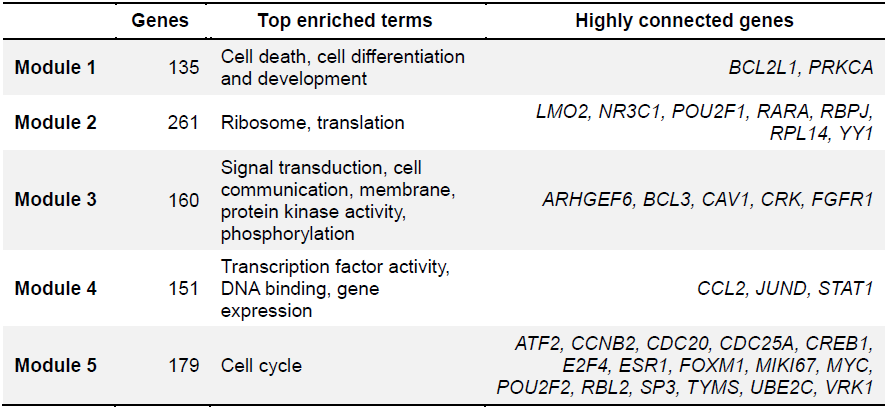
**Functional enrichment of network modules.** The first column contains the number of genes contained in every module of the network. The second column shows the top terms for which the corresponding module is enriched. The last column lists highly connected genes (degree >= 35) of the corresponding module ordered alphabetically.

The first network module is heavily enriched for cell death and apoptosis and also shows enrichments for cell differentiation, cellular development processes, and system and multicellular organismal development, as well as cancer pathways. The most connected gene in this module is *PRKCA* (Protein kinase C alpha), a kinase involved in cell differentiation, cell cycle checkpoint and cell volume control, which also plays an important role in the growth and invasion of cancers [32] and is known to act as an anti-apoptotic agent in leukemic B cells by phosphorylating *BCL2* [33]. Precisely *BCL2L1* (B-Cell CLL/Lymphoma 2 like 1), a member of the *BCL2* family, is the second most connected gene in the module.

Module two, which is enriched for the ribosome and translation as well as transcription, contains the highest connected gene of the network *POU2F1* (POU class 2 homeobox 1), a transcription factor which has been associated with the cell cycle [34] and is involved in the activation of immunoglobulin genes [35].

The signaling module (module three) in the network shows – beside heavy enrichments for signal transduction and cell communication – localization to the plasma membrane and further enrichments for kinase activity and phosphorylation. One of the highly connected genes within this module is *CAV1* (Caveolin 1), a gene strongly related to signal transduction which is able to affect cell function and cell fate [36, 37] and has furthermore been described to play a significant role in CLL progression [38]. Also, it has been shown that signaling induced by the B cell receptor *(BCR)* – which although not among the most highly connected genes is contained in network module three – in CLL cells leads to transcriptional responses of genes strongly associated with cell activation, cell cycle initiation and progression [39, 40]. It has even been suggested that part of the transcriptional differences between M-CLL and U-CLL are not cell intrinsic but secondary to *in vivo* B cell receptor stimulation [39], further emphasizing the influence of signaling and subsequent phenotypic alterations in CLL. From a technical point of view, isolation procedures activating signaling pathways through ligation of the B cell receptor could introduce a bias in these results. However, the documented relationship between single cell expression heterogeneity and plasticity of gene expression in response to perturbations [6] suggests that the primary origin of the increased response to signaling seen in U-CLL might be the higher heterogeneity present in this disease subtype.

The most connected gene in network module four, which is enriched for transcription factor activity, DNA binding and gene expression, is *JUND* (Jun D Proto-Oncogene), a member of the *AP1* transcription factor complex which regulates lymphocyte proliferation [41]. This gene has been suggested to protect cells from *p53* mediated senescence and apoptosis [42] and has an influence on tumorigenesis and cancer progression [43]. Two other highly connected genes of network module four are *CLL2* (Chemokine C-C Motif Ligand 2), a gene involved in immunoregulatory and inflammatory processes [44] which has *AP1* binding sites in its promoter [45], and *STAT1* (Signal transducer and activator of transcription 1, 91kDa), a transcriptional activator which plays an important role in lymphocyte proliferation and survival as well as cell viability in response to stimuli and pathogens [46] and has been shown to be aberrantly phosphorylated on serine residues in CLL [47]. In CLL it has furthermore been related to resistance to DNA-induced apoptosis [48].

The most important gene of the cell cycle module (module number five) is *MYC* (V-Myc avian myelocytomatosis viral oncogene homolog), a transcription factor that activates the expression of many genes but can also act as a transcriptional repressor [49]. It has a direct role in the control of DNA replication [50], drives cell proliferation and is a key player in regulating differentiation, cell growth and apoptosis by modulating the expression of distinct target genes, for example the downregulation of *BCL2* among other apoptotic pathway genes [51, 52]. Deregulated *MYC* expression has been shown to be very strongly related to tumor formation [51] and *MYC’s* expression is altered in many types of cancers [52], including CLL [53]. Further highly connected genes in the cell cycle module are *FOXM1* (Forkhead box protein M1), which plays a key role in multiple facets of cell cycle progression and is known as a proto-oncogene which contributes to both tumor initiation and progression in leukemia [54, 55] and has been shown to be upregulated in many tumors, and other key regulators of the cell cycle such as *ESR1* (Estrogen receptor 1), which is known to be involved in cell growth, cellular proliferation and differentiation [56], *RBL2* (Retinoblastoma-like 2), a progression marker gene in CLL [57], and *E2F4* (E2F transcription factor 4, p107/p130-binding), a gene which has been shown to be deregulated in rapidly growing B cell lymphomas, both of which are interacting key regulators of the cell division cycle [58], and *MKI67* (Marker of proliferation Ki-67), a gene widely used as a marker of cellular proliferation in tumors and a strong predictor of survival in CLL [59].

A possible interpretation of these data is that U-CLL patients show increased variability in proliferation rate (directly affected by the cell cycle regulation genes), cell differentiation and development, cell death, and in their intercellular communication. It is possible that U-CLL samples would show increased variability in their developmental stage, indicating the likely presence of cells at different steps of differentiation. Increased proliferation rate heterogeneity in U-CLL compared to M-CLL could be impacting this disease subtype’s aggressiveness and adaptability, possibly explaining U-CLL’s worse clinical outcome. The cell cycle status of CLL cells has been strongly related to clinical course and it has already been shown that U-CLL has more proliferative potential than M-CLL [40, 60].

### Classification of patients into the two clinical subtypes of CLL based on gene expression variability

The previous results, showing considerable differences in gene expression variability between the two clinical subtypes of CLL, suggest that gene expression variability measurements may allow the separation of M-CLL and U-CLL patients in a classification approach.

As mentioned in the studies of Kulis et al. [22] and Ferreira et al. [24], gene expression data ‘as is’ is not sufficient to cluster patients into the two classes (see also Supplementary Figure 9). Nevertheless, when applying a kind of ‘de-noising’ strategy on expression data, we are able to group the patients reasonably well into the two subtypes via unsupervised clustering (Supplementary Figure 10). This indicates that previous results of gene expression profiles not being able to distinguish the two disease subtypes are probably caused by another important aspect of gene expression variability, namely the general noise of both technical and biological origin which is present in transcriptomic data, especially at low levels of expression [26, 27].

To investigate this further, we trained a Random forest classifier [61] on the ICGC data [22] and used this classifier to predict the CLL subtypes of the patients in the Fabris et al. dataset [21]. To robustly estimate error rates, we repeated this analysis 1000 times. The classifier based on gene expression data is able to classify patients correctly (mean AUC = 0.90, see Figure 3 and Table 2).

Next, based on the promising observations we made when reducing gene expression noise, we repeated this analysis using only the top 500 most differentially expressed genes (Additional File X4) between M-CLL and U-CLL as the feature set for the Random forest classifier. The prediction of the disease subtypes in Fabris’ dataset based on the classifier trained on the ICGC data improves considerably when using the top 500 differentially expressed genes (mean AUC = 0.96, see Figure 3 and Table 2).

**Figure 3.**
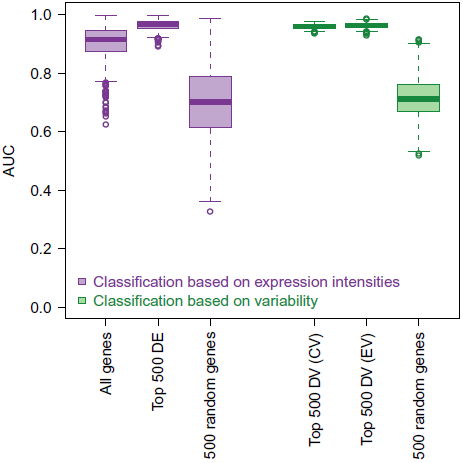
Random forest classifier results. Boxplots showing the distribution of AUC values of 1000 independent runs per classifier.

**Table 2.**
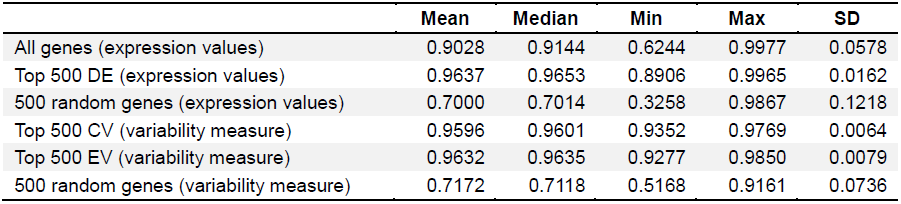
**Random forest classifier results.** AUC values of 1000 independent runs per classifier.

Finally, inspired by our results on the importance of the variability of gene expression as a defining characteristic of the two CLL subtypes, we specify a measure for each patient and each gene that could serve as a proxy for expression variability. To this end, we defined the absolute distance of a gene’s expression value from the median of expression of that gene across all patients (see Methods) and trained our Random Forest classifier applying this measure to the top 500 differentially variable genes. Again, we then aimed to predict the disease subtype of the patients in the Fabris et al. study [21] using the classifier trained on the data of Kulis et al. Strikingly, this classifier based on gene expression variability performs equally well as the one based on differential expression, with a mean AUC of 0.96 and an even smaller standard deviation, indicating more robust results compared to using mean expression levels (see Figure 3 and Table 2). Classifiers using feature sets consisting of 500 randomly selected genes perform significantly worse both in the case of using gene expression levels and the variability measure introduced above (see Figure 3 and Table 2).

In summary, as suggested by our results on differences of expression variability between the two clinical types, expression variability can classify the two subtypes remarkably well, pointing to a potentially important relation between expression variability and disease aggressiveness.

## Conclusions

We found that the more aggressive type of CLL, U-CLL, is characterized by higher variability in gene expression across patients. We additionally showed that a classifier based on gene expression variability is able to correctly classify the patients of an independent validation dataset into the two different disease subtypes, confirming the importance of expression variability in the study of CLL.

Our observation of increased variability across patients in U-CLL could be related to higher intra-patient variability in this more aggressive type of the disease, which has been observed at the genetic [18] and epigenetic [62] level. Together with these two levels of biological regulation, the contribution of drug therapy and individuals’ age [62], as well as possibly technical factors, cannot be discarded in explaining part of the observed inter-patient variability.

We showed that genes that display increased gene expression variability in the U-CLL subtype are significantly enriched for inter-cellular communication and signal transduction, which are basic components of leukemogenesis and CLL progression. Further important functions showing increased variability in U-CLL patients are related to proliferation, growth, development and apoptosis, reinforcing the possible link between increased expression heterogeneity and clinical subtypes. Actually, the combination of therapeutic agents killing cancer cells with drugs that reduce cell-to-cell variability has been suggested as a possible strategy to improve cancer treatment [64].

The observations we made in our study could also relate to single-cell heterogeneity in each patient. Currently, this hypothesis allows us to link the across-patient variability to the worse prognosis observed for U-CLL patients, which can be attributed to the presence of heterogeneity and hence aggressiveness, adaptability and resilience to drugs in the patients. To verify this hypothesis, larger datasets and single-cell genomics data would be an invaluable new source of complementary information.

## Methods

### Gene expression and methylation datasets

We used the ICGC CLL microarray datasets previously published by Kulis et al. [22] and Ferreira et al. [24]. Gene expression measurements were obtained by Affymetrix Human Genome U219 Array Plates. 48,786 features of the microarray had passed quality controls and filtering as described previously [24]. Briefly, raw CEL files were preprocessed and normalized using the RMA (Robust Multi-array Average) algorithm [65] and the Affy package [66].

The dataset comprises 122 CLL samples (70 M-CLL and 52 U-CLL) and 20 control samples of different healthy B cells (5 naive B cells, 3 IgM^+^ and IgD^+^ memory B cells, 4 IgA^+^ and IgG^+^ memory B cells and 8 CD19^+^ B cells).

DNA methylation was measured by Infinium HumanMethylation450K BeadChips. 282,470 probes (139,076 of them falling into gene promoter regions) had passed quality control and filtering procedures as described previously [22]. In summary, the data were analyzed by Genome Studio (Illumina, Inc.) and R using the lumi package [67]. To remove eventual technical and biological biases an optimized analysis pipeline was developed and applied by Kulis et al. [22]. This pipeline takes the different performance of Infinium I and Infinium II assays into account and performs additional filtering steps.

Furthermore, we included an additional gene expression dataset of CLL published by Fabris et al. [21] under GEO accession number GSE9992, containing 60 samples (24 M-CLL and 36 U-CLL) and 22,215 probes in our analyses. The microarray platform used in this study was the Affymetrix Human Genome U133A Array. The data was quality assessed and preprocessed as described previously [24]. The dataset was normalized independently from the ICGC gene expression dataset using the fRMA algorithm [68].

In order to be able to investigate the relationship between gene expression and DNA methylation, we mapped the microarray probe identifiers to Ensembl identifiers and used the average of the measurements for each gene. DNA methylation features were mapped to genomic regions (especially promoters and gene bodies) as described previously [22].

### Measuring gene expression variability

We estimated gene-wise expression variability by two different measures: i) the Coefficient of Variation (CV), defined as the ratio between the standard deviation of expression values across patients and its mean, and ii) the Expression Variability (EV) measurement proposed by Alemu et al. [27]. Alemu et al. applied local polynomial likelihood estimation [69] to model variance as a function of the mean of expression. Then the ratio of observed variance to expected variance was used as the statistic measuring expression variability, where the expected variability for each gene was estimated by a gamma regression model.

### F-test for differential variance

We performed gene-wise F-tests comparing M-CLL with U-CLL using R’s var.test function. Multiple hypotheses testing correction was performed using the Benjamini-Hochberg algorithm [70].

### Analysis of differential gene expression

Genes with differential expression between M-CLL and U-CLL were identified by limma [71]. Correction for multiple hypotheses testing was performed using the Benjamini-Hochberg algorithm [70]. Genes were considered differentially expressed when their corrected p-values are smaller than 0.05 and their absolute M-values are greater than 1.

### Creation of lists of top genes with increased variability in U-CLL

To identify the top 500 genes with increased variability in U-CLL in the dataset of Kulis et al. [22] we used genes with p-values corrected for multiple hypotheses testing smaller than 0.05 and only considered genes with consistently increased variability in U-CLL across all three variability measures employed (CV difference, EV difference, and the F-test). The remaining genes were ordered by their CV differences (CV_M-CLL_ - CV_U-CLL_) and EV differences (EV_M-CLL_ - EV_U-CLL_) respectively. In the case of the Fabris et al. dataset only 172 genes reached statistical significance, therefore we did not apply the p-value cutoff in order to achieve a comparable list of 500 genes. Both lists are available in Additional File X1.

For the list of the top 500 genes with increased variability in U-CLL in common in both datasets which is the one we used for the creation of the network (see below) we applied the same approach as described above, with the only difference that we did not cut the list after the first 500 genes within each dataset separately but when reaching 500 genes in common in both datasets. The list of these top 500 genes in common in both datasets is also available in Additional File X1.

### Functional analysis

Functional analyses were performed on the top 500 genes showing increased variability in U-CLL (available in Additional File X1). To test for enrichment of biological functions and pathways we used DAVID [72]. We uploaded the list of top 500 genes of the ICGC CLL dataset [22] and the top 500 genes of the Fabris CLL dataset [21] and used as the background set the corresponding set of genes analyzed in the dataset. We tested for the following functional annotation: GOTERM_BP_ALL, GOTERM_CC_ALL, GOTERM_MF_ALL, KEGG_PATHWAY and REACTOME_PATHWAY and set the threshold of Counts to a minimum of 3 genes. We consider terms and pathways as significantly enriched when the corresponding p-value adjusted by the Benjamini-Hochberg algorithm [70] for multiple hypotheses correction is smaller than 0.05.

### Network construction

We used the B cell specific functional interaction network of Lefebvre et al. [30] containing 5748 nodes (genes) and 64,600 edges (interactions) based on Entrez gene identifiers. We selected the 500 genes with increased variability in U-CLL in the two CLL datasets analyzed (see above) and mapped them to Entrez gene identifiers resulting in 494 unique Entrez genes. We then selected these 494 genes and their direct neighbors in the network and maintained only those genes which were at least connected with two other genes which led to a final network of 892 genes connected by 3390 edges. This network of 892 genes is the one we investigated further.

We then identified 5 network communities in the network of 892 genes by using Gephi [73] and Louvain’s method [31]. 6 genes were not mapped to any of the other network modules and were therefore excluded from the subsequent functional enrichment analyses of network modules. These enrichment analyses were performed the same way as described before, except the background gene set used which in this case is the set of all genes contained in the entire B-cell network of Lefebvre et al. [30] which are also present on the microarray platforms investigated (n = 5,548).

### Creation of feature sets for random forest classification

For establishing the feature sets used in the random forest classification approach we considered only genes present on both microarray platforms of the two different studies (Affymetrix Human Genome U219 Array Plates in the case of the data of Kulis et al. and Affymetrix Human Genome U133A Arrays in the study of Fabris et al., n = 12,307).

In order to identify the top differentially expressed genes we considered only genes with p-values corrected for multiple hypotheses testing smaller than 0.05 and ordered the genes by their absolute M-values.

For differentially variable genes we did essentially the same. We set the FDR to 5% and furthermore we only took into account genes which showed a consistently increased or decreased variability according to all three differential variability measures we applied (i.e. the CV difference, the EV difference, and the F-test) and ordered the list of the remaining genes once by their absolute CV differences (CV_M-CLL_ - CV_U-CLL_) and once by their EV differences (EV_M-CLL_ - EV_U-CLL_).

The lists of the top 500 genes used for the random forest classification can be found in Additional File X4.

### Random Forest classification

Random Forest models [61] were trained on the Kulis et al. dataset [22] (randomForest package in R, 1000 trees). Expression values were used as features for the first classifier, either all genes present on both microarray platforms used, or the top 500 differentially expressed genes (available in Additional File X4).

We further created models using the top 500 most differentially variable genes (available in Additional File X4) and defining a new feature as the distance from one gene expression value to the median of that gene over the population i.e. abs[x-median(x)]. The ROCR package was employed to calculate Area Under the Curve (AUC) values, which were used to evaluate the prediction of the Fabris et al. patients, our independent test set [21]. We ran the algorithm 1000 times independently for all classifiers in order to obtain a robust estimation of error rates.

All analyses were performed using R version 3.1.0 (x86_64-pc-linux-gnu) [74].

### List of abbreviations

AUC: Area Under the Curve
CLL: Chronic Lymphocytic Leukemia
CV: Coefficient of Variation
DV: Differential Variability
EV: Expression Variability measure of Alemu et al.
FDR: False Discovery Rate
ICGC: International Cancer Genome Consortium
ROC: Receiver Operating Characteristic

## Authors’ contribution

S. E. performed the analysis and wrote the paper. V. P. conceived the study, performed the analysis and wrote the paper. D. R. wrote the paper and guided the analysis. A. V. wrote the paper and supervised the work.

## Description of additional data files

The following additional data are available with the online version of this paper.

additional_file_1.pdf: Main supplementary file. Supplementary figures, tables and methods.

additional_file_x1.xlsx: Top 500 genes with increased variability in U-CLL. Excel table.

additional_file_x2.xlsx: Functional enrichments of genes with increased variability in U-CLL. Excel table of the results of the enrichment analysis.

additional_file_x3.xlsx: Functional enrichments of network modules. Excel table of functional enrichments of the different modules in the highly differentially variable gene network.

additional_file_x4.xlsx: Top 500 differentially variable genes. Excel table.

## Acknowledgements

This work was funded by the Spanish Ministry of Economy and Competitivity (MINECO, BIO2012-40205), the BLUEPRINT Consortium (FP7/2007-2013, under grant agreement number 282510), and the CLL Genome project (http://www.cllgenome.es) of the International Cancer Genome Consortium (ICGC), which is funded by MINECO through the Instituto de Salud Carlos III (ISCiii). The authors thank everyone in the CLL Genome project of the International Cancer Genome Consortium, in particular Elias Campos, Jose Ignacio Martin-Subero and their teams for stimulating discussions on the topic of this project. They also thank Hector Corrada Bravo for kindly providing the script to calculate the EV measurement. Furthermore, the authors gratefully thank David Juan, Victor de la Torre and all members of the Structural Biology and Biocomputing program for interesting discussions. SE is supported by a fellowship from La Caixa and VP by a FEBS long-term fellowship.

